# An accessible infrastructure for artificial intelligence using a docker-based Jupyterlab in Galaxy

**DOI:** 10.1101/2022.07.08.499333

**Authors:** Anup Kumar, Gianmauro Cuccuru, Björn Grüning, Rolf Backofen

## Abstract

Artificial intelligence (AI) programs that train on a large amount of data require powerful compute infrastructure. Jupyterlab notebook provides an excellent framework for developing AI programs but it needs to be hosted on a powerful infrastructure to enable AI programs to train on large data. An open-source, docker-based, and GPU-enabled jupyterlab notebook infrastructure has been developed that runs on the public compute infrastructure of Galaxy Europe for rapid prototyping and developing end-to-end AI projects. Using such a notebook, long-running AI model training programs can be executed remotely. Trained models, represented in a standard open neural network exchange (ONNX) format, and other resulting datasets are created in Galaxy. Other features include GPU support for faster training, git integration for version control, the option of creating and executing pipelines of notebooks, and the availability of multiple dashboards for monitoring compute resources. These features make the jupyterlab notebook highly suitable for creating and managing AI projects. A recent scientific publication that predicts infected regions of COVID-19 CT scan images is reproduced using multiple features of this notebook. In addition, colabfold, a faster implementation of alphafold2, can also be accessed in this notebook to predict the 3D structure of protein sequences. Jupyterlab notebook is accessible in two ways - first as an interactive Galaxy tool and second by running the underlying docker container. In both ways, long-running training can be executed on Galaxy’s compute infrastructure. The scripts to create the docker container are available under MIT license at https://github.com/anuprulez/ml-jupyter-notebook.

## Supplementary Note 1: Findings

### A. Background

Bioinformatics comprises many sub-fields such as single-cell, medical imaging, sequencing, proteomics and many more that produce a huge amount of biological data in myriad formats. For example, the single-cell field creates gene expression patterns for each cell that are represented as matrices of real numbers. The medical imaging field generates images of cells and tissues, radiography images such as chest x-rays and computerized tomography (CT) scans. Next-Generation sequencing generates deoxyribonucleic acid (DNA) sequences that are stored as fasta (1) files. Artificial intelligence (AI) approaches such as machine learning (ML) and deep learning (DL) have been vastly used with these datasets (2) for predictive tasks such as medical diagnosis, imputing missing datasets, augmenting datasets, and estimating gene expression patterns and many more. To be able to use ML and DL algorithms on such datasets, a robust and efficient compute infrastructure is needed that can serve multiple purposes. They include pre-processing raw datasets to transform them into suitable formats that are understood by ML and DL algorithms, creating and executing complex architectures of ML and DL algorithms on preprocessed datasets and making models and predicted datasets readily available for further analyses. To facilitate such tasks, a complete infrastructure is developed that combines jupyterlab (3) notebook, augmented with many useful features, running on public compute resources of Galaxy (4) Europe to perform end-to-end AI analysis on biological datasets. The infrastructure consists of three important components. First, a docker container that encapsulates jupyterlab together with packages such as git (5), elyra AI (6), tensorflow GPU (7), CUDA (8) for tensorflow to interact with GPU and many others. Second, a Galaxy interactive tool (9, 10) downloads this docker container to serve jupyterlab online on Galaxy Europe. Third, the compute infrastructure, consisting of several CPUs and GPUs, on which the online jupyterlab runs.

### B. Jupyterlab

Jupyterlab is a web-based, robust editor used for varied purposes such as data science, scientific computing, machine learning and deep learning. It’s a common editor that supports more than 40 programming languages, some of the popular ones are python, R, julia and scala. Python is one of the most popular languages used by researchers for performing numerous scientific and predictive analyses. Therefore, it is used as the programming language in Galaxy jupyterlab interactive tool because many popular packages such as tensorflow, pytorch (11), scikit-learn (12) for machine learning and deep learning are readily available as python packages. Moreover, the extensible architecture of jupyterlab makes it possible to add many external packages and plugins such as git, dashboards and many others. Editors such as jupyterlab, integrated with several useful packages, provide a favourable platform for both, rapid prototyping and end-to-end development and management of AI projects. To harness the benefits of jupyterlab, it has been used as the editor for an interactive tool in Galaxy.

### C. Docker container

Docker (13) containers are popular for shipping packaged software as complete ecosystems enabling them to be reproducible in a platform-independent manner. Software executing inside a docker container is abstracted from the operating system (OS) as most of the requirements necessary for them to run successfully are already configured inside the container. A container runs as an isolated environment making the minimum number of interactions to the host OS and thereby, improving the security aspects of running software. Docker container is used in this project to encapsulate jupyterlab along with many useful packages such as CUDA, tensorflow, ONNX (14), scikit-learn and many others. Docker container inherits many packages such as numpy (15), scipy (16) from its base container, Jupyter/tensorflow-notebook (17), and augments it with many other packages suitable for machine learning, deep learning and visualisation. Docker container is independent of Galaxy and can be separately executed for serving jupyterlab with the same set of packages on a different compute infrastructure or any personal computer (PC) and laptop. Moreover, it can easily be extended by installing suitable packages only by adding their appropriate package names in its dockerfile (18). The docker container then should be rebuilt and uploaded to the docker hub (19). Galaxy interactive tool can automatically download the updated container from the docker hub and the newly installed packages become available in the jupyterlab notebook.

### D. Features

Many features such as easy accessibility, support of a wide variety of programming languages of jupyterlab, and extensibility to install useful plugins make it a desirable editor for researchers, especially for creating prototypes rapidly. In this project, many such features have been integrated into a jupyterlab notebook which is served online on Galaxy Europe with large compute resources to enable researchers to create prototypes and end-to-end AI projects (Figure 1). Some of the important features of jupyterlab hosted on Galaxy Europe are GPU support for faster execution of deep learning programs. Tensorflow-GPU interacts with GPU using CUDA when the backend compute resource has GPU(s), otherwise, the program in a jupyterlab notebook runs on CPUs. Other useful packages include ONNX for transforming trained tensorflow and scikit-learn models to ONNX models, open-CV (20) and scikit-image (21) for processing images, nibabel (22) for reading image files stored as “.nii”, bioblend (23) for connecting to Galaxy to access its datasets, histories and workflows in a jupyterlab notebook and visualisation package such as bqplot (24) for plotting interactive charts, voila (25) for displaying output cells of a jupyterlab notebook, dashboard such as nvdashboard (26) for monitoring GPU performance. Support for the file extension such as H5 (27), efficient for storing matrices, enables machine learning researchers to save model weights and input datasets for an AI algorithm. Other packages such as colabfold (28) together with JAX (29) are used for predicting 3D structures of proteins which are discussed in detail in a later section. In addition, it is possible to create a long-running training job that runs on a remote compute infrastructure and the trained models and output datasets get stored permanently in Galaxy history. The trained model is saved as an ONNX file and tabular datasets in H5 file.

**Fig. 1.**
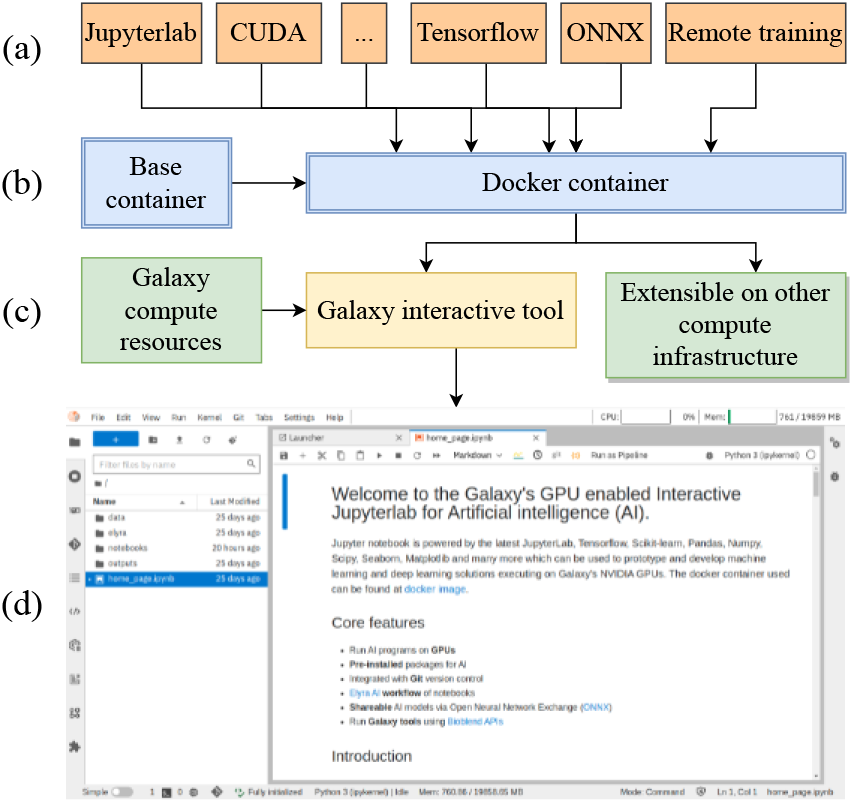
Architecture of Galaxy’s jupyterlab. Part (a) shows packages and features wrapped inside a docker container. Part (b) shows a base docker container (17) from which the customized container (19) is derived. In part (c) Galaxy’s interactive tool downloads the customized container. The customized docker container can also be hosted on a different compute infrastructure. Part (d) shows Galaxy’s jupyterlab.

### E. Related infrastructure

There are a few infrastructures available, free and commercial, that offer jupyterlab or similar environments for developing data science and AI projects. A few popular ones are google colab (30), kaggle kernels (31) and amazon’s sagemaker (32). Google colab is partially free and offers an online editor similar to a jupyterlab notebook. The free version of colab offers dynamic compute resources. The disk space is around 70 gigabytes (GB) and memory (RAM) is around 12 GB. These resources are scarce for projects that deal with high-dimensional biological data (33, 34). These resources are also variable and depend on past usage. More compute resources are assigned to those users that have used less in the past for a more equitable sharing of resources. Moreover, there is a limit of the running time of only 12 hours which is also insufficient and not ideal for long-running model learning training on large datasets. However, colab pro and pro+ offer better compute resources but they come at a price - EUR 9.25 and EUR 42.25 per month, respectively. In contrast, kaggle kernels are free of charge but similar to colab, their computing resources are scarce. The total disk space is approximately 73 GB and RAM is 16 GB for a CPU-based kernel. For the GPU-based kernel, the disk space is of the same size as that of the CPU-based kernel but the RAM of the CPU decreases to 13 GB and an additional RAM of 15 GB is added through GPU and computation time is limited to 30 hours a week. It also supports TPUs but the computation time is even more limited to only 20 hours a week. Amazon’s sagemaker is also a commercial software for developing AI algorithms that is free of charge but only for 2 months. Overall, these notebook infrastructures do not offer unrestricted compute resources free of charge. To address the drawbacks of these notebook servers and provide researchers and users large compute resources more reliably, Galaxy jupyterlab offers 1 terabyte (TB) of disk space and unlimited computation time on 1 GPU and 7 CPUs per session. RAM for GPU is around 15 GB and for CPUs is 20 GB (Table 1). The offered resources in the jupyterlab notebook running in Galaxy stay constant and are independent of past usage. To more it more useful, jupyterlab opens a tab for each notebook that allows researchers to develop and execute all notebooks inside the same session of the allotted computational resource rather than forcing them to connect to a different session as in google’s colab and kaggle kernels.

**Table 1.**
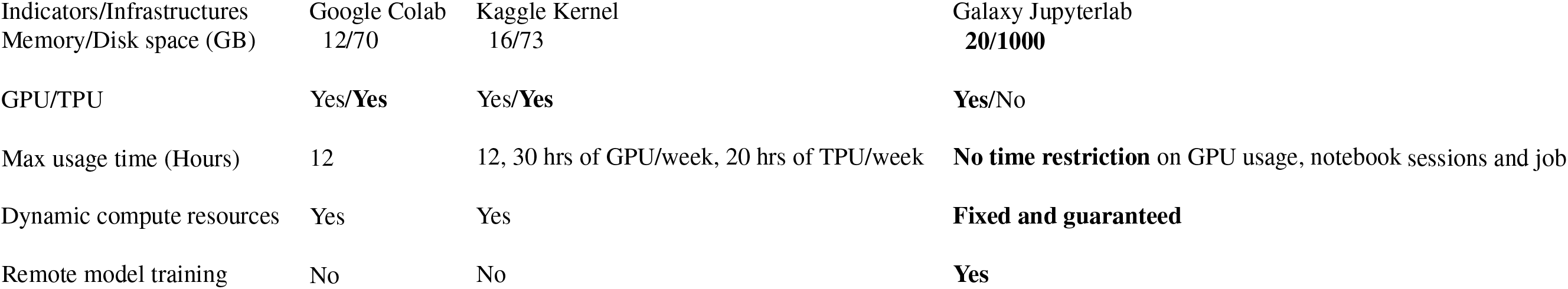
Comparison with other notebook infrastructures

## Supplementary Note 2: Implementation

Jupyterlab infrastructure has been developed in two stages. First, a docker container is created containing all the necessary packages such as jupyterlab itself, CUDA for interaction with GPU, tensorflow, scikit-learn, ONNX and so on. The docker container is inherited from a base container that is suited for serving entire jupyterlab environments. In addition to software packages such as numpy, scipy, and tensorflow that are already wrapped around the base container, many packages are added with their compatible versions. Compatible packages for CUDA, CUDA DNN and tensorflow are necessary so that they together interact with the GPU on the host machine for accelerating deep learning programs. Other significant packages, integrated into the docker container, are ONNX, scikit-learn, elyra AI, bioblend, nibabel, scikitimage, open-CV, bqplot and voila. The integrated docker container contains all the necessary packages for developing data science, machine learning and deep learning projects. Second, the container can be downloaded to any powerful compute infrastructure and jupyterlab can be served in a browser via the URL it creates. In addition, to run this container in Galaxy, an interactive tool is created that downloads this container on a remote compute infrastructure and generates a URL that is used to run jupyterlab in a browser. The architecture of jupyterlab infrastructure in Galaxy is shown in Figure 1.

The running instance of Jupyterlab in Galaxy contains a home page, a jupyterlab notebook, that summarises several of its features. Further, there are other notebooks available, each describing a feature of the jupyterlab with code examples such as how to create ONNX models for scikit-learn and tensorflow classifiers, how to connect to Galaxy using bioblend, how to create interactive plots using bqplots and how to create a pipeline of notebooks using elyra AI. To access the jupyterlab notebook in Galaxy Europe, a ready-to-use hands-on GTN (35) tutorial (36) has been developed that shows steps such as opening the notebook, using git to clone codebase from github, and sending long-running training jobs to a remote Galaxy cluster. The two use-cases, explained in previous sections, are also covered in the tutorial along with their respective code scripts as separate jupyterlab notebooks.

### .1. Remote model training

For large datasets, model training may need several hours or even days. In such cases, it would be non-ideal to keep the jupyterlab notebook open in a browser’s tab till the training finishes. Therefore, another Galaxy tool (37) is developed to enable researchers to send long-running training jobs to a remote Galaxy cluster. The tool can be executed from a jupyterlab notebook using a custom python function (38) that takes input datasets and training script as input parameters. The input datasets to be used for training, testing and validation must be provide in H5 format. This is done to standardise input data format for AI models that train on matrices as input data can be in multiple formats such as images, genomic sequences, gene expression patterns. H5 files can be created using any of these data formats and fed to the AI model. The long-running training happens in a remote cluster in Galaxy as a regular job. Upon completion of the job, the resulting datasets and the trained model become available in a newly created Galaxy history. This feature of outsourcing deep learning’s long-running training to a remote cluster decouples it from the jupyterlab notebook. The trained model and other datasets can be downloaded from the Galaxy history for further analysis.

## Supplementary Note 3: Results

Jupyterlab infrastructure in Galaxy is used to reproduce two scientific publications that demonstrate its power to develop deep learning models using COVID CT scan images (39) and predict the 3D structure of proteins using colabfold, a faster implementation of alphafold2 (40).

### A. COVID-19 CT scan image segmentation

In (39), COVID-19 CT scan images have been used to develop and train a deep learning model architecture that predicts COVID-19 infected regions in CT scan images with high accuracy. An open-source implementation of the work is available that trains a unet deep learning architecture that distinguishes between normal and infected regions in CT scans. The code of this implementation is adapted and executed on our jupyterlab notebook infrastructure. The CT scan images used in the original work (39) are transformed into an H5 file so that they can directly be used as an input to the unet architecture defined in the jupyterlab notebook (41). A composite H5 file (42) is created using script (43) that contains multiple datasets inside and each dataset is a matrix corresponding to the matrices training, test and validation as used in the original work. The entire analysis of the original work can be reproduced using multiple notebooks in (41) by achieving similar values of precision and recall (approximately 0.98) as mentioned in the original work. In (41), the first notebook (1_fetch_datasets.ipynb) downloads input dataset as an H5 file. Additionally, it also downloads the trained ONNX model. The second notebook (2_create_model_and_train.ipynb) creates and trains a unet model on the training dataset extracted from the H5 file. Training, accelerated by GPU, for 10 iterations over the entire training dataset finishes quickly. The third notebook (3_predict_masks.ipynb) extracts the test dataset and predicts infected regions of the CT scans in the test dataset using the trained model created by the second notebook. Figure 2 shows the comparison of ground truth infected regions (second column) and the predicted infected regions in the third and fourth columns. Original CT scans from the test dataset are shown in the first column of Figure 2.

**Fig. 2.**
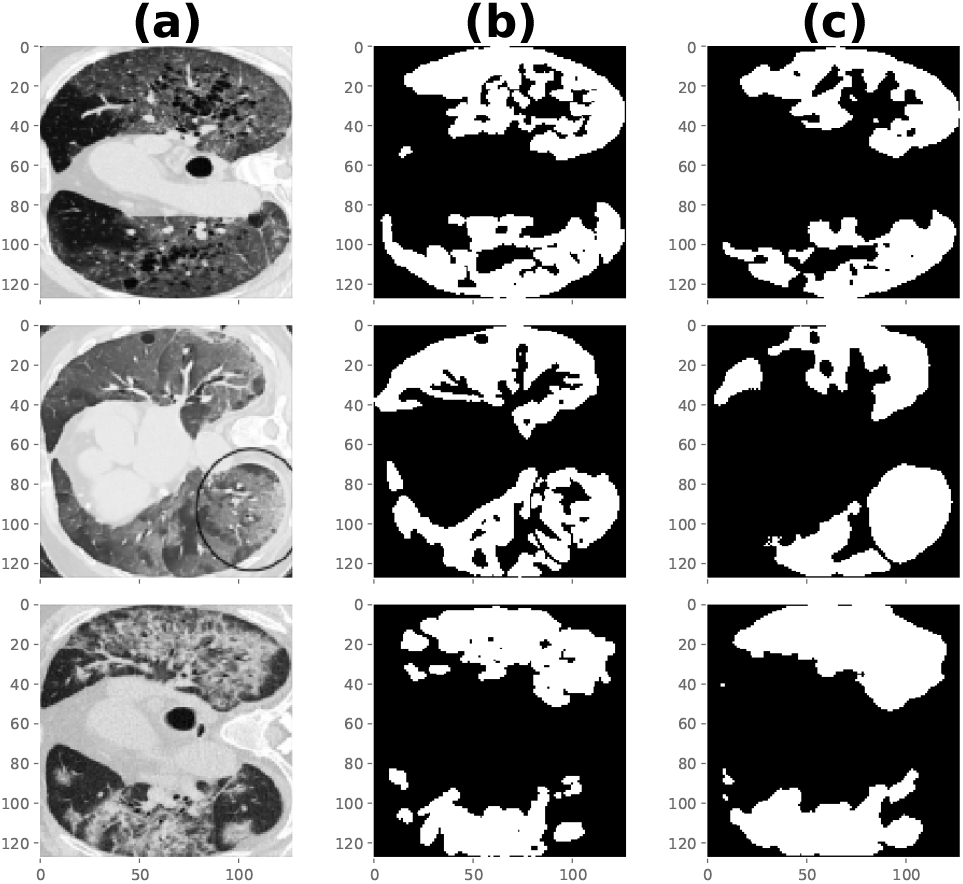
Figures shows original CT scans in column (a), corresponding ground-truth masks of original CT scans in column (b) and the predicted masks in column (c). Masks are COVID-19 infected regions in the corresponding CT scans. The groundtruth and predicted masks show high similarity.

In (41), the notebook “4_create_model_and_train_remote.ipynb” defines the entire script for developing and training the unet architecture. “5_run_remote_training.ipynb” notebook executes the previous notebook on a cluster remotely after creating a Galaxy history and then uploading the script extracted from “4_create_model_and_train_remote.ipynb” notebook and datasets. The custom python function (run_script_job) creates a Galaxy history using bioblend and then uploads the datasets to the same history. After the upload is finished, the python script from the specified notebook is executed dynamically. It trains a deep learning model on the uploaded datasets to create a model which is saved as an ONNX model in Galaxy history. Using the notebook “6_predict_masks_remote_model.ipynb” from (41), the trained model can be downloaded from the Galaxy history and used for predicting infected regions of the CT scans of the test dataset. A significant advantage of training deep learning models remotely is that researchers don’t have to keep the jupyterlab notebook session running as long as the model is being trained. It provides a convenient way for training models, especially those that take several hours or even days to finish.

### A.1. Predict 3D structure of proteins using ColabFold

Al-phafold2 has made a breakthrough in predicting the 3D structure of proteins with outstanding accuracy. However, due to their large database size (a few TB), it is not easily accessible to researchers. Therefore, a few approaches have been developed that replace the time-consuming steps of alphafold2 with slightly different steps but predict the 3D structure of proteins with similar accuracy while consuming less memory and time. One such approach is colabfold which replaces a large database search in alphafold2 for finding homologous sequences by a significantly (40-60 times) faster MM-seqs2 API (44) call to generate input features based on the query protein sequence. Colabfold’s prediction of 3D structures in batches is approximately 90 times faster. Colabfold is integrated into the docker container (19) by adding two packages - colabfold and GPU-enabled JAX which is a justin-time compiler for making mathematical transformations. “7_ColabFold_MMseq2.ipynb” notebook in (41) makes the prediction of 3D structure using colabfold by making use of the alphafold2 pre-trained weights. Figure 3 shows the 3D structure of 4Oxalocrotonate_Tautomerase (45), a protein sequence of length 62, along with its side chains. This 3D structure is extremely similar to the structure predicted by the jupyter notebook (46) from colabfold in (28).

**Fig. 3.**
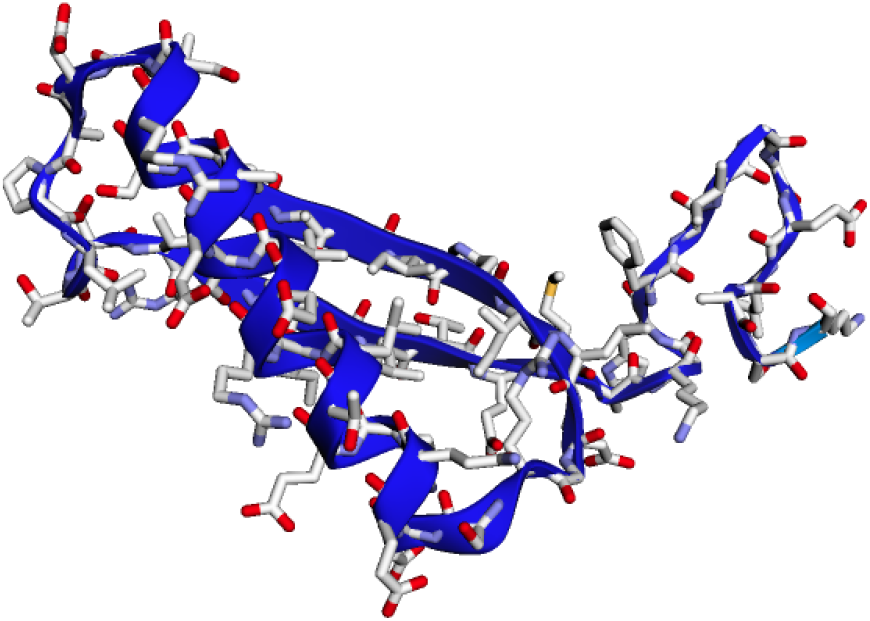
Figures shows a 3D structure of 4 Oxalocrotonate_Tautomerase enzyme (protein) predicted by ColabFold

## Supplementary Note 4: Summary

Jupyterlab notebook is integrated as an interactive tool in Galaxy Europe running on a powerful compute infrastructure comprising CPUs and GPUs. Jupyterlab is configured in a docker container along with many different packages such as CUDA, tensorflow, scikit-learn, and elyra AI to provide a robust architecture for the development and management of projects from data science, machine learning and deep learning. Remote training makes it convenient to run multiple analyses in parallel as different Galaxy jobs by executing the same Galaxy tool and results become available in different Galaxy histories. Features such as git integration are useful for managing entire code repositories on github and elyra AI for creating pipelines of notebooks to be executed as one unit of software. All notebooks run on the same session of the jupyterlab. The entire infrastructure of jupyterlab is readily accessible through Galaxy Europe. In contrast to commercial infrastructures that host editors similar to Jupyterlab and offer powerful and reliable compute only through paid subscriptions, this infrastructure provides large compute resources invariant to usage and has unlimited usage time while ensuring the same set compute resources for multiple usages.

## Supplementary Note 5: Availability of supporting source code and requirements

Project name: GPU-enabled docker container with Jupyterlab for artificial intelligence

Project home page: https://github.com/anuprulez/ml-jupyter-notebook

Galaxy interactive tool: https://github.com/usegalaxy-eu/galaxy/blob/release_22.01_europe/tools/interactive/interactivetool_ml_jupyter_notebook.xml

Operating system: Linux

Programming languages: Python, XML, Docker, Bash

Other requirements:

License: MIT License

Biotools ID: gpu-enabled_docker_container_with_jupyterlab_for_ai

## Supplementary Note 6: Declarations

## A. List of abbreviations

GB: Gigabyte
AI: Artificial intelligence

## Author Approvals

All authors have seen and approved the manuscript, and it hasn’t been accepted or published elsewhere.

## C. Authors’ contributions

Authors’ contributions follow the order of names.

## D. Acknowledgements

We thank Galaxy Europe for its support.

